# Uncovering the mechanisms of real-world attentional control over the course of primary education

**DOI:** 10.1101/2020.10.20.342758

**Authors:** Nora Turoman, Ruxandra I. Tivadar, Chrysa Retsa, Anne M. Maillard, Gaia Scerif, Pawel J. Matusz

## Abstract

Schooling may shape children’s abilities to control their attention, but it is unclear if this impact extends from control over visual objects to encompass multisensory objects, which are more typical of everyday environments. We compared children across three primary school grades ([country] 1^st^, 3^rd^, and 5^th^ grade) on their performance on a game-like audio-visual attentional control task, while recording their EEG. Behavioral markers of *visual* attentional control were present from 3^rd^ grade (after 2 years of schooling), whereas multisensory attentional control was not detected in any group. However, multivariate whole-brain EEG analyses (‘electrical neuroimaging’) revealed stable patterns of brain activity that indexed both types of attentional control – visual control in all groups, and multisensory attentional control from 3^rd^ grade onwards. Multivariate EEG approaches can uncover otherwise undetectable mechanisms of attentional control over visual and multisensory objects and characterize how they differ at different educational stages.

**Lay Abstract:** We measured how visual and audiovisual distractors differ in capturing the attention of 1^st^- to 5^th^-graders while recording the children’s brain activity. Brain activity results showed that all children were sensitive to visual distraction, and from 3^rd^ grade onwards, children were also sensitive to audiovisual distraction. These results deepen our understanding of how school children control their attention in everyday environments, which are made up of information that stimulates multiple senses at a time.

---

The start of school marks a transition from a less regulated, play-oriented environment to one with increasing demands for focused attention and ignoring distractors. The development of such attentional control and its relationship with schooling experience remains poorly understood, especially in real-world environments.

Most current knowledge about children’s attentional control has come from research on executive functions (EF; Miyake et al., 2000), which are closely linked with attentional control (Bavelier & Green, 2019). Both EF’s and attentional control gradually improve over childhood (e.g., Donnelly et al., 2007) due to protracted structural changes within and between the prefrontal and parietal cortex (e.g., Casey et al., 2005). Such findings have clarified when and how children’s control skills compare to those of adults, rather than helped map how control skills function during various stages of childhood, when children are challenged by their educational environment in different ways. Though both approaches are important, we focus on the latter for two reasons.

First, the development of control skills need not be uniformly linear when the multisensory nature of the environment and the child’s schooling experience are considered. Matusz and colleagues (2015) found that while 11-year-olds and adults were distracted by audiovisual shape-sound stimuli in a visual search task, 6-year-olds were immune to such distraction. Thus, young children’s limited attentional control can shield them from real-world distraction, rather than making more distracted than adults, as visual-attentional research typically suggests. Similar developmental differences were observed for distraction by digits (Matusz et al., 2019a). Children with less schooling experience (i.e., less familiar with numerals) were more protected from distraction by conjunctions of visual numerals and their auditorily-presented names. Second, children’s attentional control skills are linked to scholastic success. Schooling is a catalyst for developing cognitive control, e.g., IQ increases with education level (e.g., Brinch & Galloway, 2012), and EF skills improve when children enter formal schooling (e.g., Brod et al., 2017). Yet, the influence of schooling on attentional control in multisensory contexts is currently unknown.

To support education better we need to understand how attention is deployed in multisensory environments like classrooms. Multisensory processes undergo development, alike visual-attentional processes. Already infant brains are sensitive to congruency across the senses (e.g., Lewkowicz & Turkewitz, 1980), but processes integrating weighted inputs mature much later (+8 years; e.g., Gori et al., 2008). Studying attentional control and *distraction* gauged by audio-visual objects impacts classroom learning. Distraction by unisensory content hinders classroom learning (vision: Godwin & Fisher, 2011; hearing: Massonnié et al., 2019). Skills in multisensory letter-sound mappings predict scholastic achievements (Bach et al., 2013) similarly to visual EF/attentional control skills (e.g., Bull et al., 2008).

Here, we pursued two questions: 1) How do children process multisensory vis-à-vis visual distractors, and what brain mechanisms govern these processes? 2) How audio-visual control changes with school experience? We investigated the behavioral and brain mechanisms of attentional control using a child-friendly multisensory spatial-cueing paradigm while recording EEG. We conducted traditional (N2pc) and multivariate (electrical neuroimaging) analyses of the event-related EEG potentials (ERPs). We expected older children to show visual attentional control behaviorally; we had no strong hypotheses for multisensory attentional control or the underlying EEG mechanisms.

## Methods

### Participants

In [country], children enter formal education at age 4, where the first two years are considered kindergarten. By 3^rd^ grade (ages 6-7) children sit at desks and receive more structured classroom instruction. We tested 92 children from local primary schools: 26 5^th^-graders (10 male, *M±SD_age_* 8y10mo*±*5mo, range: 8y1mo–10y1mo), 38 3^rd^-graders (18 female, *M±SD_age_* 6y10mo*±*4mo, range: 6y1mo–7y9mo), and 28 1^st^-graders (13 female, *M±SD_age_* 5y*±* 4mo, range: 4y–5y7mo; full details in Supplementary Online Materials, SOMs). All research procedures were approved by the local ethics committee.

### Stimuli and procedure

Participants were tested in the local hospital in an experimental session lasting 1h–1h30mins, where we recorded their performance and EEG. The paradigm was a multisensory variant of Folk et al.’s (1992) spatial cueing paradigm (Matusz & Eimer, 2011, Exp.2) adapted to be child-friendly (Figure 1).

**Figure 1.**
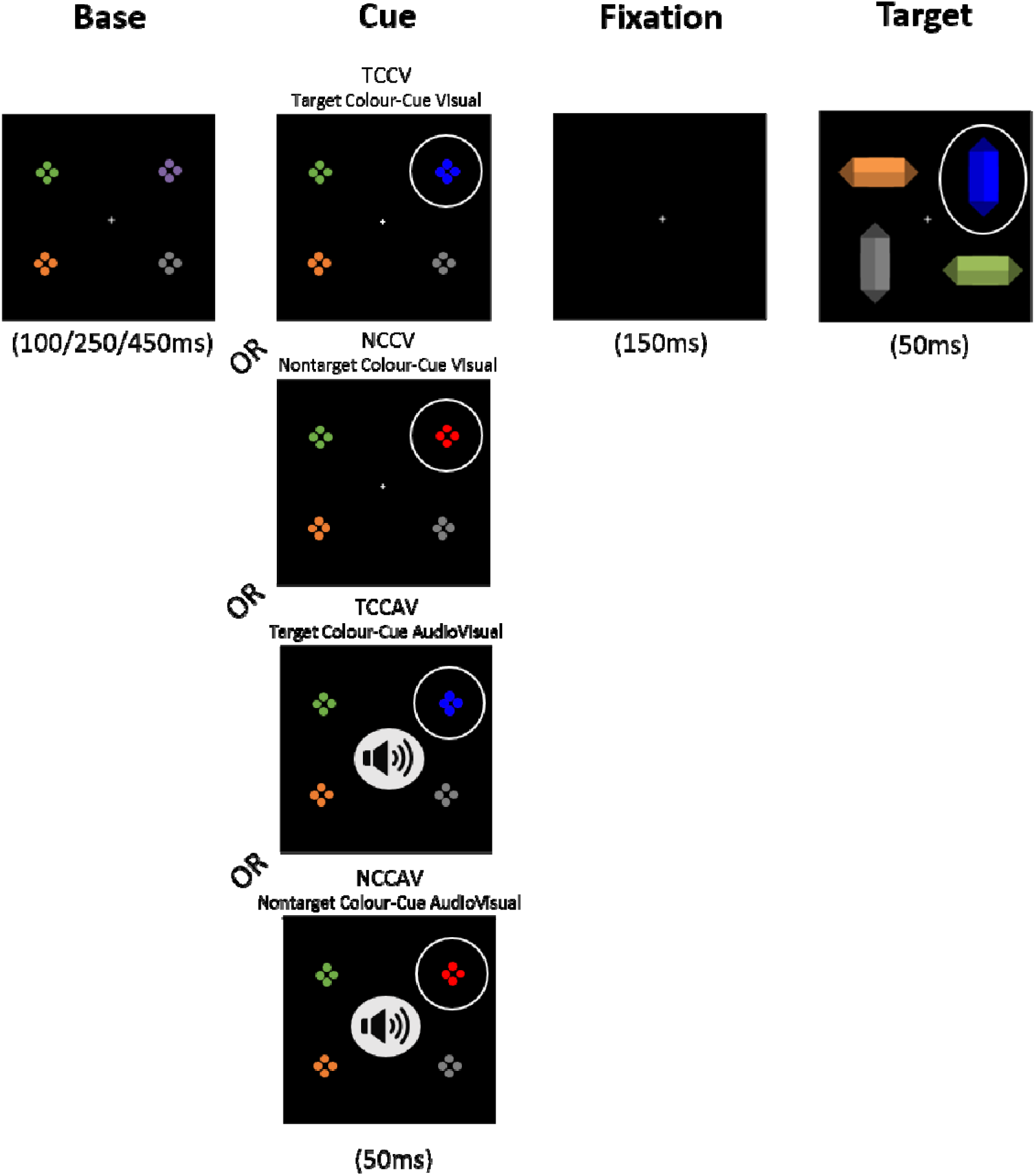
Experimental trial sequence for our paradigm. Base array contained four differently colored elements. In the cue array, one of these elements changed color – to a target-matching color (blue cue for a blue diamond), or a nontarget color (red cue). On 50% all trials, the color-change cue was accompanied by a tone (indicated symbolically by a speaker). Cue color and sound presence manipulations created the four cue conditions (shown in figure). Cue and target appeared in the same (indicated symbolically by white circles) or different location.

Participants searched for a target diamond of a predefined color, and responded as quickly and accurately as possible to the target’s orientation (horizontal or vertical; randomly determined on each trial) by pressing one of two large buttons (Lib Switch, Liberator Ltd.) fixed onto a tray on their lap. Each diamond was preceded by a cue, which matched the target color (e.g., blue for blue target) or did not (red for blue target; red and blue were counterbalanced as target/cue colors across participants). Location of the cue was randomized and unpredictive of the location of the target, thus eliciting attentional capture. On 50% of trials, the cue coincided with the onset of a pure sine-wave tone (2000Hz). These manipulations reflect the two cue factors in our design: Cue Color (target-color cue vs. nontarget-color cue) and Cue Modality (visual vs. audiovisual), producing 4 cue conditions: target-color cue visual, nontarget-color cue visual, target-color cue audiovisal, target-color cue audiovisal (Figure 1). We measured these effects of visual and multisensory attentional control through our main dependent variable: difference in raw speed between trials where the cue and the target shared their location (thus the cue captured attention, this way facilitating response speed) versus trials where the target appeared in a location different from the preceding cue’s (after the cue captured attention, attention had to be disengaged from the cue location and onto the target location, leading to slower responses). The third design factor, Cue-Target Location, defined by this difference between speeding and slowing (Same vs. Different location) of raw responses, measured how strongly a given cue captured attention. See SOMs for further details of the procedures.

### EEG acquisition and preprocessing

Continuous EEG was recorded using a 129-channel HydroCel Geodesic Sensor Net (1000Hz sampling rate) connected to a NetStation amplifier (Net Amps 400; Electrical Geodesics Inc.). Impedances were kept <50kΩ, and electrodes online referenced to Cz. Preprocessing involved: offline filtering (0.1 Hz high-pass, 40 Hz low-pass, 50 Hz notch, and a second-order Butterworth filter with a linear –12 dB/octave roll-off with forward and backward passes to eliminate phase-shift), segmentation into epochs around cue onset (−100ms; 500ms), semi-automated rejection of transient noise, eye movements, and muscle artefacts. Artefact rejection criteria were ±150 μV, with visual inspection (Shimi et al., 2014). Artefact-contaminated electrodes were interpolated using three-dimensional splines (average numbers of epochs removed and interpolated electrodes, in SOMs). Cleaned epochs were averaged, baseline-corrected (100ms pre-cue time-interval), and re-referenced to the average reference. EEG/ERP analyses were anchored to the cue array. An additional 50Hz notch filter was applied due to persistent environmental noise despite initial filtering. Only data from trials with correct responses and from blocks with >50% accuracy were analyzed.

Preprocessing was done separately for ERPs from the four cue conditions, and for cues in left and right hemifields. To analyze cue-elicited lateralized ERPs, data from both hemiscalps was anchored to a ‘reference’ side. Labels of single-trial data from trials with cues presented on the left were relabelled to represent activity over the right hemiscalp, creating veridical “cue-on-the-right” data and mirrored “cue-on-the-right” data. Next, we averaged these two data types, for the four cue conditions, creating four single lateralized average ERPs. We performed all preprocessing and analyses using Cartool (www.fbmlab.com/cartool-software/).

### Data analysis design

#### Behavioral analyses

We analyzed mean reaction-time (RT) attentional capture effects, following related literature (Folk et al., 1992; Gaspelin et al., 2015; RT cleaning described in SOMs). We submitted mean RTs to a mixed 4-way repeated-measures ANOVA (rmANOVA) with a between-subject factor Age (5^th^-graders vs. 3^rd^-graders vs. 1^st^-graders) and three within-subject factors: Cue Color (target-color cue vs. nontarget-color cue), Cue Modality (visual vs. audiovisual), and Cue-Target Location (same vs. different). Here, task-set contingent attentional capture was tested via a Cue-Target Location × Cue Color interaction (**stronger** attentional capture by target-color cues but not nontarget-color cues). Multisensory enhancement of attentional capture was assessed via a Cue-Target Location × Cue Modality interaction (stronger attentional capture for audiovisual than visual cues). All analyses were conducted using SPSS. Detailed behavioral results were reported in our previous study on adult-like audio-visual attentional skills over childhood (reference post-review), so here we report only the most relevant results.

EEG analyses. The N2pc component is a spatially-selective enhancement of negative potentials over occipital electrodes on the side contralateral (vs. ipsilateral) to the selected object. It is a traditional marker of attentional selection (Eimer 1996). We combined N2pc analyses with an electrical neuroimaging (EN) approach. As no reliable cue-elicited N2pc’s were found in any age group, we report only the EN analyses of the cue-elicited ERPs in the N2pc time-window (see SOMs for details of the N2pc analysis design and results).

The EN approach encompasses a set of multivariate, reference-independent analyses of global features of the electric field at the scalp across the whole electrode montage. Combining EN measures with established ERP correlates of cognitive processes can help elucidate the cognitive and brain mechanisms underlying multisensory attentional control (e.g., Matusz et al. 2019b), as EN analyses are capable of brain mechanisms that give rise to modulations in ERPs by visual and multisensory control. To obtain global EN measures of *lateralized* N2pc effects, the contralateral-ipsilateral difference ERPs created for N2pc analyses were mirrored onto the other hemiscalp, constructing ‘mirrored’ 129-channel datasets. From these ‘mirrored’ 129-channel difference-ERPs, Global Field Power (GFP) and topographical EEG patterns were analyzed.

For GFP analyses, each age group’s mean GFP over their N2pc time-windows were extracted from group-averaged ‘fake’ ERPs per condition and submitted to separate two-way rmANOVAs with factors: Cue Color and Cue Modality. Differences in the GFP would indicate that visual and/or multisensory control (main effects of Cue Color and Cue Modality, respectively) modulate cue-elicited lateralized ERPs by altering the *strength* of response within a similar (statistically indistinguishable) brain network (detailed explanations of GFP and topographic analyses for lateralized ERPs in SOMs, also Matusz et al. 2019b; Turoman et al. 2020).

For topographical analyses, first, stable periods of topographic activity (“topographic maps”) were identified through a clustering (“segmentation”) procedure, which was conducted on group-averaged ERPs. Here, we clustered the ERPs for the four cue conditions within each grade’s respective N2pc time-window. The optimal set of maps was chosen based on largest global variance they explain, and the cross-validation and Krzanowski-Lai criterions. Clustering necessarily means that similar patterns have been identified across one participant group (here, grade). Then, the results of the segmentation of the grand-averaged ERPs are fit back onto the single-subject data to see how much each of the maps identified in the segmentation characterizes individual participants. This is how we obtained map durations (in milliseconds) over each child’ N2pc time-window, which we then submitted to three separate three-way rmANOVAs, with factors: Cue Color, Cue Modality, and Map (different levels due to different numbers of maps in each age group). Differences in topographic maps would indicate that visual and/or multisensory control (Map × Cue Color and Map × Cue Modality interactions, respectively) modulated cue-elicited lateralized ERPs by altering the recruited brain networks. Multiple comparisons between map durations were Holm-Bonferroni corrected. Comparisons passed the correction unless otherwise stated.

## Results

### Behavioral analyses

A main effect of Age, *F*_(2, 89)_=32.8, *p*<0.001, η_p_^2^=0.4 revealed that mean RTs sped up reliably from 1^st^-graders (1309ms) through 3^rd^-graders (1107ms) to 5^th^-graders (836ms; all *p*’s<0.001, see SOMs). Although Age did not interact with other factors (all *F’s*<2, *p*’s>0.1), RT capture effects were analyzed per age group to clarify multisensory distraction effects across school grades.

Fifth-graders showed visual task-set contingent attentional capture (Cue-Target Location × Cue Color interaction, *F*_(1,25)_=19.5, *p*<0.001, η_p_^2^=0.4); their attention was captured by target-color cues (56ms), but not nontarget-color cues (Figure 2A). However, their attention was not enhanced for audiovisual than visual cues (no Cue-Target Location × Cue Modality interaction, *F*_(1,25)_=1.4, *p*=0.3; Figure 2A). Third-graders also showed task-set contingent attentional capture (*F*=6.4, *p*=0.02, η_p_^2^=0.2), with their attention captured by target-color cues (55ms), but not nontarget color-cues (Figure 2B). Third-graders also showed no evidence for multisensory enhancement of attentional capture (*F*_(1,37)_=2.1, *p*=0.2; Figure 2B). Contrastingly, first-graders showed no evidence for visual task-set contingent attentional capture (*F*_(1,27)_=1.4, *p*=0.2) or multisensory enhancement of attentional capture (*F*_(1, 27)_=0.4, *p*=0.5; Figure 2C).

**Figure 2.**
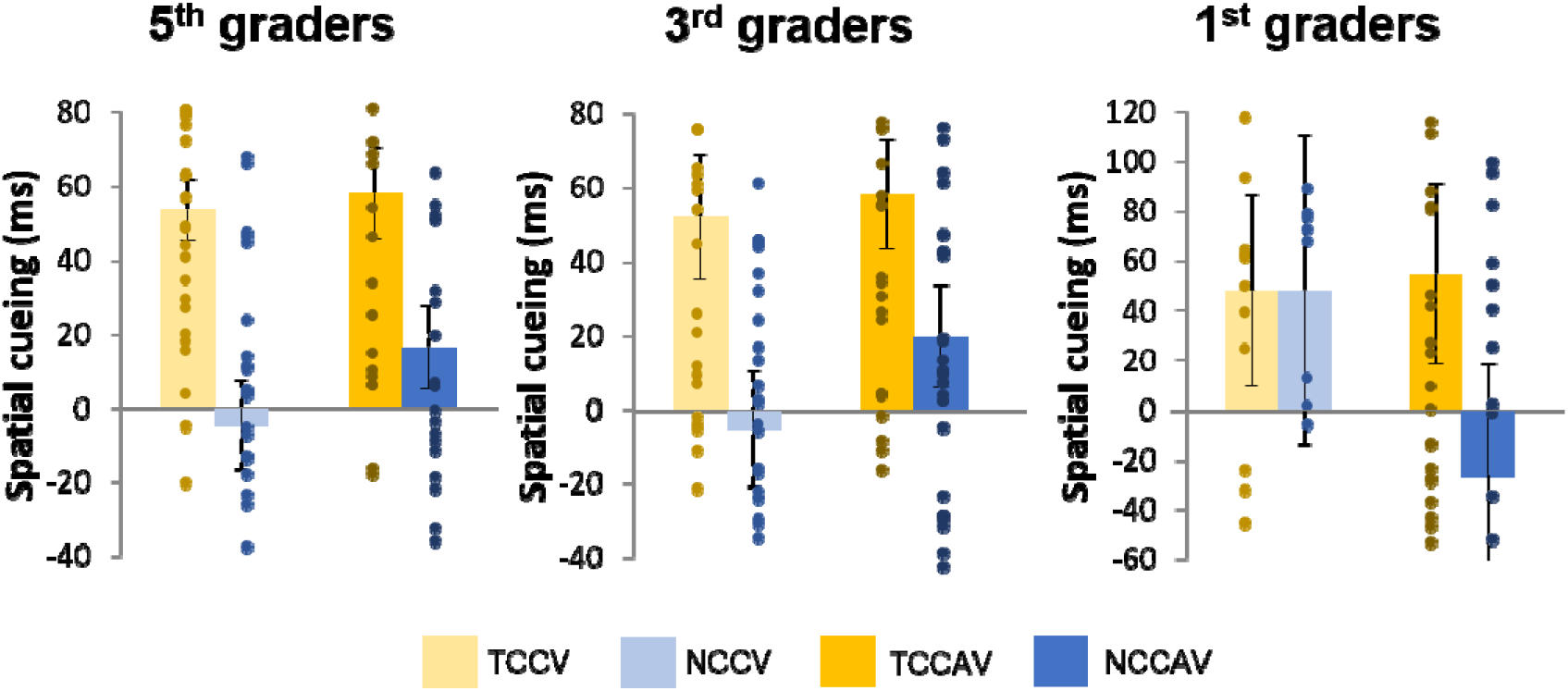
Mean RT attentional capture effects. Error bars represent standard errors of mean (S.E.M.’s), and vertical dots - individual RT attentional capture effects. Third- and 5^th^-graders showed larger behavioural capture for target-color than nontarget-color cues; no groups showed multisensory capture enhancements.

### EEG analyses

Three separate 2 × 2 rmANOVAs on the average GFP over each age group’s N2pc time-window revealed no main effects or interactions in 5^th^- and 1^st^-graders (*p*’s>0.1), with a trend in 3^rd^-graders for a main effect of Cue Color (*F*=3.07, *p*=0.09). Full results are reported in SOMs.

Segmentations of the post-cue period per age group resulted in: 14 clusters in 5^th^-graders (explaining 91.5% of global variance in the group-averaged difference ERPs), 11 clusters in 3^rd^-graders (88.3% global-explained variance), and 11 clusters in 1^st^-graders (84.9% global-explained variance). Next, the fitting procedure revealed the template maps that characterized each age-group’s N2pc time-window: 5^th^-graders - 9 maps over 144–290ms; 3^rd^-graders - 5 maps over 151–275ms; 1^st^-graders - 8 maps over 110–302ms post-cue. Age-specific maps were differentiated using grade-related prefixes (‘5’ for 5^th^-graders, etc.). Statistically non-significant map duration differences (*p*’s>0.1) were not reported.

In 5^th^-graders, a 2 × 2 × 9 rmANOVA revealed a main effect of Map in ERPs within N2pc time-window, *F*=2.7, *p*=0.009, η_p_^2^=0.1, confirming they show stable patterns of topographic lateralized activity (that are captured by N2pc). Follow-up analyses focused on comparisons investigating visual and multisensory attentional control in ERP topography. Visual control modulated the topography of 5^th^-graders’ cue-elicited ERPs (Map × Cue Color interaction, *F*_(8,200)_=3.4, *p*=0.001, η_p_^2^=0.1), which was driven by three maps: Map54, Map56, and Map59 (Figure 3A, left panel). Map56 was more active during processing of target-color cues than nontarget-color cues (28 vs. 8ms, *t*_(25)_=3.7, *p*=0.005). Meanwhile, Map54 and Map59 were more active during processing of nontarget-color cues than target-color cues (Map54 20 vs. 7ms, *t*_(25)_=3.4, *p*=0.008; Map59 16ms vs. 9ms, *t*_(25)_=3, *p*=0.04,Figure 3A, left panel).

**Figure 3.**
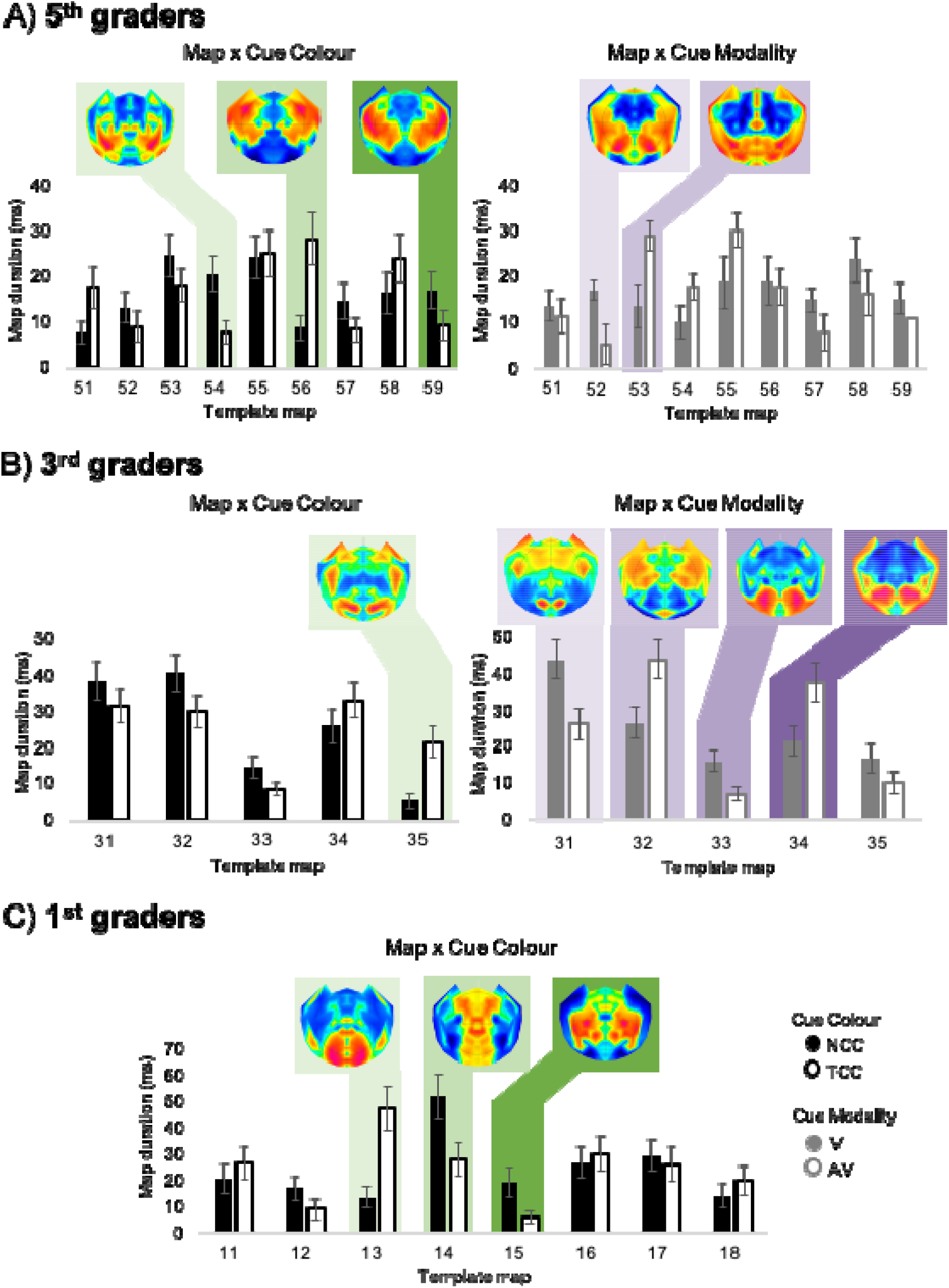
EN analyses of visual and multisensory control effects in ERP topography. Bars represent mean durations (ms) of maps present in N2pc time-window in each age group, where black/black-outlined bars represent Map × Cue Color interaction, grey/grey-outlined bars - Map × Cue Modality interaction. Maps in shades of green and purple were modulated Cue Color and Cue Modality, respectively. Error bars represent S.E.M.’s.

Also multisensory control modulated topography of 5^th^-graders’ ERPs (Map × Cue Modality interaction, *F*_(8,200)_=2.4, *p*=0.02, η_p_^2^=0.1), driven by two maps: Map52 and Map53 (Figure 3A, right panel). Map53 was more active during processing of audiovisual than visual cues (29ms vs. 13ms, *t*_(25)_=3.7, *p*=0.007), while Map52 was more active during processing of visual than audiovisual cues (17ms vs. 5ms, *t*_(25)_=3.1, *p*=0.03). Visual and multisensory control also interacted (Map × Cue Color × Cue Modality interaction, *F*_(8,200)_=2.2, *p*=0.048, η*_p_²*=0.1; post-hoc analyses in SOMs).

In 3^rd^-graders, a 2 × 2 × 5 rmANOVA revealed a main effect of Map, *F*_(3.2,117.6)_=9.8, *p*<0.001, η_p_^2^=0.2. Visual control modulated topography of 3^rd^-graders’ cue-elicited ERPs (Map × Cue Color interaction, *F*_(2.9,107)_=2.8, *p*=0.04, η_p_^2^=0.1), driven by Map35 being more active during processing of target-color cues than nontarget-color cues (21ms vs. 5ms, *t*_(37)_=3.7, *p*=0.002, Figure 3B, left panel). Third-graders’ ERP topography was modulated by multisensory control (Map × Cue Modality interaction, *F*_(3.1,114.3)_=8, *p*<0.001, η_p_^2^=0.2), driven by Maps 31–34 (Figure 3B, right panel). Specifically, Map32 and Map34 were more active during processing of audiovisual over visual cues (Map32 44ms vs. 26ms, *t*_(37)_=3.5, *p*=0.002; Map34 37ms vs. 21ms, *t*_(37)_=3.2, *p*=0.01). Conversely, Map31 and Map33 were more active during processing of visual over audiovisual cues (Map31 43ms vs. 26ms, *t*_(37)_=3.5, *p*=0.004; Map33 16ms vs. 6ms, *t*_(37)_=2.8, *p*=0.006). Again, visual and multisensory control interacted (Map × Cue Color × Cue Modality interaction, *F*=3.2, *p*=0.03, η_p_*²*=0.1; post-hoc analyses in SOMs).

In 1^st^-graders, a 2 × 2 × 8 rmANOVA revealed a main effect of Map, *F*_(4.7, 127.2)_=4, *p*=0.003, η_p_^2^=0.1. Like in the older groups, visual control modulated 1^st^-graders’ cue-elicited ERP topography (Map × Cue Color interaction, *F*_(7,189)_=4.2, *p*<0.001, η_p_^2^=0.1), driven by Maps 13–15. Map13 was more active during processing of target color-cues over nontarget color-cues (47ms vs. 13ms, *t*_(27)_=4.3, *p*=0.003). Conversely, Map14 and Map15 were more active during processing of nontarget-color cues than target-color cues (Map14 52ms vs. 27ms, *t*_(27)_=3.1, *p*=0.007; Map15 19ms vs. 5ms, *t*_(27)_=2.8, *p*=0.05). Unlike the older groups, however, 1^st^-graders showed no evidence for multisensory control in ERP topography (no Map × Cue Modality, interaction, *F*_(7,189)_=0.9, *p*=0.4). That said, visual and multisensory control interacted (Map × Cue Color × Cue Modality, *F*_(7,189)_=2.2, *p*=0.04, η_p_^2^=0.1; post-hoc analyses in SOMs).

## Discussion

Using multivariate analyses of the N2pc, a well-known ERP marker of attentional selection in adults, we have shown that brains of children are sensitive to visual task-relevance of objects and objects’ multisensory nature early during primary education.

### Top-down visual attentional control is present even at school entry

In behavior, robust feature-specific (color) goal-based visual attentional control, indexed by task-set contingent attentional capture, was observed in 3^rd^- and 5^th^-graders. This is younger than what most extant research on attentional control processes demonstrates. Importantly, EN topographical analyses demonstrated distinct stable patterns of global brain activity that were sensitive to such visual control already at school entry.

In older groups, EN revealed the brain mechanisms underlying the behaviorally-observed patterns of visual attentional control. In 3^rd^-graders, behavioral task-set contingent attentional capture may emerge from enhanced target-matching distractor (target-color cue) processing, via recruitment of brain networks preferentially processing goal-relevant information. In 5^th^-graders, task-set contingent attentional capture may be driven by a combination of enhanced processing of goal-relevant information and suppressed processing of goal-irrelevant (nontarget-color cue) information: One map was primarily active during the processing of target-matching distractors, two other maps – of nontarget-matching distractors. While it cannot be ascertained if increased presence of nontarget-matching maps shows the inhibition of goal-irrelevant information, the concomitant behavioral inhibition of nontarget-matching distractors would support this notion. A similar pattern of results was found in adults’ visual N2pc’s (Hickey et al., 2009). Though the relationship between topographic map modulations and distractor processing is not clear-cut, we reveal that children govern visual feature-based (color) attentional control via differential brain network recruitment (not in brain response strength, i.e., gain control).

In 1^st^-graders, EN revealed nascent visual attentional control. Despite no behavioral task-set contingent attentional capture, 1^st^-graders showed two maps predominating responses to nontarget-matching distractors, and one map recruited for target-matching distractors, mirroring our findings in 5^th^-graders. These results strengthen past findings by directly showing early onset of feature-specific top-down visual control at 4 years and adding novel mechanistic insights at the brain level. Most studies on control processes in 4-year-olds used behavioral (e.g., Bull et al., 2008; Gaspelin et al. 2015) or heamodynamic measures (e.g., Brod et al., 2017; Fiske & Holmboe 2019). It is a novel, exciting finding that separable sets of brain networks are preferentially active in response to goal-relevant and goal-irrelevant information even earlier than the 5-to-7-year shift.

### Attentional control over multisensory objects develops after two years of schooling

Behavioral analyses did not detect multisensory enhancement of capture in any group. However, in the older groups, EN revealed distinct brain networks over 200ms post-stimulus recruited by visual *and* audiovisual distractors. This finding supports the idea that salient multisensory stimuli, even when task-irrelevant, control attention independently of top-down goal-relevance (Matusz & Eimer, 2011). Our results suggest that multisensory distraction emerges earlier than previously thought. While previous studies demonstrated that multisensory interference develops only around 6-7 years (Matusz et al., 2015; 2019a), we show that involuntary attention to multisensory objects develops already after two years of schooling. Thus, 6-7-year-olds are not protected from it, as previous work would suggest.

First-graders showed no evidence for multisensory attentional control modulating cue-elicited ERP topography. If anything, sounds accompanying visual distractors attenuated their visually-elicited ERPs, as shown by suppressed contralateral ERP responses. It may be that in 1^st^-graders, but not older children, attentional resources are separably allocated to vision and audition (e.g., Welch & Warren, 1980, but see Matusz et al. 2015). We are currently investigating how children at school-entry process the additional distractor sounds.

Our EN analyses revealed that <1 year of schooling experience affords children’s brain networks the sensitivity to goal-relevance of visual stimuli; 2 additional years - to audiovisual stimuli. Our EN results were mirrored by behavioral results (partly, for visual control) but not N2pc results. By extension, the EN results uncovered despite null behavioral results (for multisensory control; potentially driven by still-developing motor processes, e.g., Kail & Ferrer, 2007) should also be genuine effects. The validity and reproducibility of the identified topographic maps is supported by several direct sources of evidence. First, their optimal number was selected based on criteria of residual noise, reliability, and optimal map configuration. Second, the maps were statistically analyzed at single-subject level, during the fitting procedure. Finally, maps derived across similar tasks are highly reproducible, both in clinics (Baradits et al. 2020) and basic research. For example, different groups across 10 studies reliably identify 4-7 EEG resting-state maps, which match the networks identified using MRI (Michel & Koenig 2018).

### Studying developing attentional control with EEG/ERPs

Crucially, our findings reveal EN analyses as more sensitive than canonical N2pc analyses. Why? First, mounting evidence suggests that the N2pc is not an automatic marker of attentional selection, readily measurable whenever attention is studied with ERPs. It might require optimal conditions to appear, spanning characteristics related to participant (age), attended stimuli (physical, e.g., bright; cognitive, e.g., task-relevant features) and/or the task (no other stimuli within 200-250ms after presentation of main stimulus). We discuss those points in detail in the Supplemental Discussion. Second, this lower sensitivity is well explained by the ability of multivariate methods to capture patterns in data that univariate analyses are not sensitive to, e.g., spatial regularities in brain activity across time and/or experimental conditions (e.g., Matusz et al. 2018; Kriegeskorte et al. 2007).

This issue is especially relevant for neurophysiological data. Recording from only one electrode of the EEG montage forcibly reduces the sensitivity of traditional ERP analyses, by decreasing the amount of signal, and through the experimenter’s decision that activity at other electrode sites is irrelevant to the studied cognitive process: for N2pc - the control of attention in space (we discuss the limitations this brings for neuroscience and psychology in Matusz et al. 2019b). Also, by relying on the choice of a reference electrode, results from canonical ERP analyses are necessarily less reproducible, across time, labs, etc. Finally, by recording from the same electrodes across *all* participants, those analyses rely on a fundamental albeit hard to defend assumption - that brain anatomy of all participants is uniform.

EN analyses surpass all those limitations; they are independent of the reference electrode, are data-driven, and consider data from the whole electrode montage. Importantly, EN analyses are robust also against the variability in the underlying brain anatomy – if one set of brain sources in a participant is too different from the whole sample, a given map will simply not be fit to them, but their remaining brain activity will be captured by other maps. This makes EN analyses also more reproducible (more details in Supplemental Discussion). Together, our approach, involving comparing specific cognitive processes, systematically across children from specific age groups (and adults; Turoman et al. 2021), and analyses of well-known ERP correlates with multivariate approaches, holds multiple advantages for investigating the development of cognitive and brain mechanisms of attentional control.

### Implications for education

Our findings confirm that schooling supports neurocognitive development and enrich them with ideas on real-world distraction. Effects of in-classroom clutter and noise (+5 years; Fisher et al., 2014; Massonnié et al., 2019) may be exacerbated by children’s sensitivity to distraction by audiovisual objects, but only from age 6. Thus, classrooms, but not kindergartens, could support learning by reducing decoration or use of new technologies – unless these are related to the subject of instruction.

## Acknowledgments

We thank The EEG Brain Mapping Core of the Center for Biomedical Imaging (CIBM) for providing the infrastructure. We thank Noémie Kirscher for assistance with child IQ data collection and Louise Vasa for assistance with child EEG data analysis. We would also like to thank Micah Murray for helpful comments during study design and manuscript preparation. Finally, we would like to thank all of the families who participated in the current study. This work was supported by the Pierre Mercier Foundation and the Swiss National Science Foundation (PZ00P1_174150) to PJM.

## References

1. Bach, S., Richardson, U., Brandeis, D., Martin, E., & Brem, S. (2013). Print-specific multimodal brain activation in kindergarten improves prediction of reading skills in second grade. Neuroimage, 82, 605–615.

2. Baradits, M., Bitter, I., & Czobor, P. (2020). Multivariate patterns of EEG microstate parameters and their role in the discrimination of patients with schizophrenia from healthy controls. Psychiatry Research, 288, 112938.

3. Bavelier, D., & Green, C. S. (2019). Enhancing Attentional Control: Lessons from Action Video Games. Neuron, 104(1), 147–163.

4. Brinch, C. N., & Galloway, T. A. (2012). Schooling in adolescence raises IQ scores. Proceedings of the National Academy of Sciences, 109(2), 425–430.

5. Brod, G., Bunge, S. A., & Shing, Y. L. (2017). Does one year of schooling improve children’s cognitive control and alter associated brain activation?. Psychological Science, 28(7), 967–978.

6. Bull, R., Espy, K. A., & Wiebe, S. A. (2008). Short-term memory, working memory, and executive functioning in preschoolers: Longitudinal predictors of mathematical achievement at age 7 years. Developmental Neuropsychology, 33(3), 205–228.

7. Casey, B. J., Tottenham, N., Liston, C., & Durston, S. (2005). Imaging the developing brain: what have we learned about cognitive development?. Trends in Cognitive Sciences, 9(3), 104–110.

8. Donnelly, N., Cave, K., Greenway, R., Hadwin, J. A., Stevenson, J., & Sonuga-Barke, E. (2007). Visual search in children and adults: Top-down and bottom-up mechanisms. Quarterly Journal of Experimental Psychology, 60(1), 120–136.

9. Eimer, M. (1996). The N2pc component as an indicator of attentional selectivity. Electroencephalography and Clinical Neurophysiology, 99(3), 225–234.

10. Fiske, A., & Holmboe, K. (2019). Neural substrates of early executive function development. Developmental Review, 52, 42–62.

11. Folk, C. L., Remington, R. W., & Johnston, J. C. (1992). Involuntary covert orienting is contingent on attentional control settings. Journal of Experimental Psychology: Human perception and performance, 18(4), 1030.

12. Gaspelin, N., Margett-Jordan, T., & Ruthruff, E. (2015). Susceptible to distraction: Children lack top-down control over spatial attention capture. Psychonomic Bulletin & Review, 22(2), 461–468.

13. Godwin, K., & Fisher, A. (2011). Allocation of attention in classroom environments: consequences for learning. Proceedings of the Annual Meeting of the Cognitive Science Society, 33(33).

14. Gori, M., Del Viva, M., Sandini, G., & Burr, D. C. (2008). Young children do not integrate visual and haptic form information. Current Biology, 18(9), 694–698.

15. Hickey, C., Di Lollo, V., & McDonald, J. J. (2009). Electrophysiological indices of target and distractor processing in visual search. Journal of Cognitive Neuroscience, 21(4), 760–775.

16. Kail, R. V., & Ferrer, E. (2007). Processing speed in childhood and adolescence: Longitudinal models for examining developmental change. Child Development, 78(6), 1760–1770.

17. Kriegeskorte, N., Formisano, E., Sorger, B., & Goebel, R. (2007). Individual faces elicit distinct response patterns in human anterior temporal cortex. Proceedings of the National Academy of Sciences, 104(51), 20600–20605.

18. Lewkowicz, D. J., & Turkewitz, G. (1980). Cross-modal equivalence in early infancy: Auditory–visual intensity matching. Developmental Psychology, 16(6), 597.

19. Massonnié, J., Rogers, C. J., Mareschal, D., & Kirkham, N. Z. (2019). Is classroom noise always bad for children? Frontiers in Psychology, 10, 381.

20. Matusz, P. J., & Eimer, M. (2011). Multisensory enhancement of attentional capture in visual search. Psychonomic Bulletin & Review, 18(5), 904.

21. Matusz, P. J., Broadbent, H., Ferrari, J., Forrest, B., Merkley, R., & Scerif, G. (2015). Multi-modal distraction: Insights from children’s limited attention. Cognition, 136, 156–165.

22. Matusz, P. J., Key, A. P., Gogliotti, S., Pearson, J., Auld, M. L., Murray, M. M., & Maitre, N. L. (2018). Somatosensory plasticity in pediatric cerebral palsy following constraint-induced movement therapy. Neural Plasticity, 2018.

23. Matusz, P. J., Merkley, R., Faure, M., & Scerif, G. (2019a). Expert attention: Attentional allocation depends on the differential development of multisensory number representations. Cognition, 186, 171–177.

24. Matusz, P. J., Turoman, N., Tivadar, R. I., Retsa, C., & Murray, M. M. (2019b). Brain and cognitive mechanisms of top–down attentional control in a multisensory world: Benefits of electrical neuroimaging. Journal of Cognitive Neuroscience, 31(3), 412–430.

25. Michel, C. M., & Koenig, T. (2018). EEG microstates as a tool for studying the temporal dynamics of whole-brain neuronal networks: a review. Neuroimage, 180, 577–593.

26. Miyake, A., Friedman, N. P., Emerson, M. J., Witzki, A. H., Howerter, A., & Wager, T. D. (2000). The unity and diversity of executive functions and their contributions to complex “frontal lobe” tasks: A latent variable analysis. Cognitive Psychology, 41(1), 49–100.

27. Shimi, A., Kuo, B. C., Astle, D. E., Nobre, A. C., & Scerif, G. (2014). Age group and individual differences in attentional orienting dissociate neural mechanisms of encoding and maintenance in visual STM. Journal of Cognitive Neuroscience, 26(4), 864–877.

28. Turoman, N., Tivadar, R., Retsa, C., Murray, M. M., & Matusz, P. J. (2020). How we pay attention in naturalistic settings. BioRxiv.

29. Welch, R. B., & Warren, D. H. (1980). Immediate perceptual response to intersensory discrepancy. Psychological Bulletin, 88, 638–667.

