## Supplementary Online Materials for "Uncovering the mechanisms of real-world attentional control over the course of primary education"

Turoman, Tivadar, Retsa, Maillard, Scerif, Matusz

**Supplementary Methods**

**Participants**

A total of 115 primary school children were recruited for the study, 28 of whom were enrolled in 5^th^-grade, 46 in 3^rd^-grade, and 41 in 1^st^-grade. The above participants were part of a larger sample of children that were recruited from March 2017 to May 2019. Children were recruited from local schools, nurseries, public events and entertainment facilities. No children had an FSIQ under 85 (assessed using Full Scale IQ items from the WISC-V for 5th graders and 3rd graders, and the WPPSI-IV for 1st graders), which would warrant exclusion form the study. All participants had normal or corrected-to-normal vision and normal hearing, and had no history of sensory, neurological, or learning difficulties, as indicated by parental report. Informed consent was obtained from parents/caregivers and verbal assent was obtained from children before participating in the study. 18 children were excluded for failure to initiate the testing session or failure to complete the task with above chance-level accuracy (50%), thus excluding 1 5^th^-grader, 6 3^rd^-graders, and 11 1^st^-graders, respectively. Finally, 5 additional participants (1 5^th^-grader, 2 3^rd^-graders, and 2 1^st^-graders) were excluded for unusable EEG signals due to excessive noise even after the two-step filtering process.

**Stimuli and Procedure**

The task was adapted to children by reducing the number of elements per stimulus array from six to four. To increase engagement, we introduced a game-like narrative where participants had to help a pirate captain find treasure on a deserted island, and, accordingly, reshaped target bars into “diamonds”.

All trials were presented on a 23” LCD monitor with a resolution of 1080 × 1024 (60-Hz refresh rate, HP EliteDisplay E232) at 90cm viewing distance. Base array duration was randomized between the three durations shown on the figure, in order to avoid possible entrainment effects. Base array elements were comprised of closely aligned dots, subtending 0.1° × 0.1° of visual angle. Colors of elements that were not the cue or target could be one of the following, according to the RGB scale: green (0/179/0), pink (168/51/166), gold (150/134/10), and silver (136/136/132). Cue and target colors were either blue (RGB values: 31/118/220) or red (RGB values: 224/71/52), and target color was counterbalanced across participants. The diamond-like shape of the targets was achieved by adding triangle shapes on the short sides of rectangular bar shapes and increasing and decreasing the luminance of certain sides of the bars by 20%. All elements were approximately equiluminant (~18 cd/m2) and spread equidistally along the circumference of an imaginary circle against a black background, at an angular distance of 2.1° from a central fixation point. Tones presented on 50% of trials had an intensity of 80 dB SPL, as measured next to participants’ ears, and were played from two loudspeakers on the left and right sides of the stimulus display monitor.

A training block of 32 trials at 50% of regular task speed was administered before the experimental task. The experimental task was initiated once a score above chance-level accuracy (50% of maximum accuracy) was achieved. The experimental task consisted of 8 blocks of 64 trials each (total of 512 trials). During the task, participants were required to respond within a maximum of 5000ms of the target presentation. The next trial was automatically presented after a button press, or after the maximum response time has elapsed. Feedback on accuracy was given after each block, followed by a treasure map image which informed participants of the number of blocks remaining until the end of the task (i.e., reaching the treasure). When the image was displayed, participants could take a break, parents/caregivers could enter the testing room, and stickers on diamond-shaped sticker sheets were offered to maintain motivation in younger participants. As compensation for participating in the experiment, children received a 30 Swiss franc voucher for a media store, and parents/caregivers’ travel costs were reimbursed.

**EEG preprocessing**

Average numbers of epochs removed were as follows: 11% in 5th graders, 8% in 3rd graders, and 14% in 1st graders. Average numbers of electrodes interpolated in each age group were as follows: 11 in 5th graders (8% of the total electrode montage), 10 in 3rd graders (8% of the total electrode montage), and 12 in 1st graders (10% of the total electrode montage).

**N2pc data analysis design**

The N2pc is a negative-going brain response observed around 200ms post-stimulus over posterior electrodes contralateral to stimulus location (Luck & Hillyard, 1994; Eimer, 1996), which has been observed in response to target-matching distractors (e.g., Eimer et al., 2009; Eimer & Kiss, 2008). In the present study, contralateral and ipsilateral average ERPs elicited across the 4 cue conditions were first analyzed within an N2pc framework, with a departure from the canonical N2pc literature. A canonical N2pc analysis typically assumes extracting mean voltage amplitudes from a pair of posterior electrodes contralateral and ipsilateral to stimulus presentation (PO7/8) over ~200ms post-cue (e.g., Eimer, 1996; Eimer & Kiss, 2008; Eimer et al., 2009; Kiss et al., 2008, Luck & Hillyard, 1994). However, these electrode sites and time-window were validated on adult participants. Children’s N2pc’s (to targets) show different amplitudes and/or onset latencies (Couperus & Quirk, 2015; Sun et al. 2018; Shimi et al. 2015), likely due to the known physiological differences between children’s and adults’ skulls and brains (e.g., Scerif et al., 2006). We thus probed each age group’s ERP data for the electrode sites and time-window in which the N2pc would be best detected.

We first computed a contralateral-ipsilateral difference ERP by subtracting the voltage amplitudes over the contralateral hemifield (left side of the scalp) from voltage amplitudes over the ipsilateral hemifield (right side of the scalp) for the experimental condition which best resembled the stimulus settings in which the N2pc was traditionally observed, i.e., the target color matching visual condition – target-color cue visual distractor. Next, to obtain N2pc time-windows, for each group’s grand-averaged difference ERP for target-color cue visual distractors, we identified the post-cue time-period at which the DISS measure did not change for at least 120ms, assuming a typical N2pc duration. The resulting time windows were as follows: 144-271ms (127ms duration) in 5^th^ graders, 151-275ms (125ms duration) in 3^rd^ graders, and 110-310ms (192ms duration) in 1^st^ graders. Next, to identify the appropriate electrode sites for each group, in each group’s grand-averaged ERP for the target-color cue visual distractor, we identified the locations on the scalp with the highest negative voltage amplitudes over each group’s respective N2pc time-windows using an automatic setting in the CarTool software. The resulting contralateral-ipsilateral electrode sites were as follows: e68/e94 in 5^th^ graders, e68/e94 in 3^rd^ graders, and e50/e101 in 1^st^ graders. Mean amplitude values extracted from each group’s respective electrode sites and time-window were submitted to separate three-way repeated measures ANOVAs, with factors: Cue Color (Target-color cue vs. Nontarget-color cue), Cue Modality (Visual vs. Audiovisual), and Contralaterality (Contralateral vs. Ipsilateral).

**EN data analysis design**

In the electrical neuroimaging (EN) framework, differences in the strength of a response within a nonidentified brain generator are measured by Global Field Power (GFP), which is a standard deviation of voltage potential [μV] across the entire electrode montage. Differences in activation of distinct configurations of brain generators are measured by Global Dissimilarity (DISS), which is directly related to the spatial correlation of signals across two conditions.

Modulations of N2pc by goal-relevance are traditionally assumed to arise from a modulation of response strength, i.e., a “gain-control” mechanism, which EN can detect as GFP differences between target-color cue and nontarget-color conditions over the N2pc time-window (see Matusz et al., 2019). However, in contrast with the classical gain-control account, Matusz et al. (2019) suggest that differences in N2pc amplitudes between experimental conditions can be supported by different sets of globally distributed networks being active for different conditions. As part of EN, differential recruitment of brain generators underlying ERPs can be revealed by analyzing the topography of the ERPs in question.

Briefly, applying data clustering methods onto DISS over the time course of an ERP can reveal periods of tens to hundreds of milliseconds of stable topographic activity, i.e., topographic “maps”, within that ERP (elsewhere referred to as “functional microstates”, e.g., Michel & Koenig, 2018). Here, we used the Topographic Atomize and Agglomerate Hierarchical Clustering (TAAHC) approach, which identifies sequences of topographical maps in the group-averaged ERP data across all conditions. Initially, each map in the dataset belongs to one cluster, but over a series of iterations, the number of clusters becomes progressively smaller. Each remaining cluster is assigned groups of maps whose mathematical mean (centroid) represents a ‘template map’ that labels that cluster. This iterative clustering (“segmentation”) procedure generates configurations of topographical map clusters, each accounting for a portion of the global explained variance (GEV) in the group-averaged ERP data. On each iteration, the cluster with the smallest GEV is identified, dissolved, and freed of its maps, which then get reassigned to new clusters. The ‘free’ maps are reassigned to the cluster whose centroid they are best spatially correlated with (i.e., that have the highest Pearson correlation). This procedure ends with the entire dataset characterised by one cluster, which is not informative. The optimal number of clusters is based on the: maximal global explained variance (GEV) in the group-averaged ERP data with the smallest number of template maps. This number can be identified using a range of criteria, and here, we used the Cross-Validation index – a measure of residual noise (so the lower the criterion, the better) – and the modified Krzanowski–Lai criterion - a measure related to the distance between cluster members (Murray et al., 2008). During the ‘fitting’ procedure, each mirrored-global ERP datapoint over the respective age group’s N2pc time-window was labelled by the template map – from the optimal configuration of clusters for that age group revealed by group-averaged segmentation - present over their N2pc time-window with which it best correlated spatially. The resulting template map durations (in milliseconds) were submitted to analyses, as described in the manuscript. Maps which with durations under 10 contiguous timeframes were not included in the analyses. Greenhouse-Geiser corrections were applied where necessary to correct for violations of sphericity. Unless otherwise stated in the results, map durations were statistically different from 0ms (as confirmed by post-hoc *t*-tests), meaning that they were reliably present across the time-windows of interest.

**Supplemental behavioural analysis design**

Cleaning of RT data involved the following: 1) removing blocks with accuracy below chance level (50%). Overall, 15% of all blocks were removed (5^th^-graders: 3%, 3^rd^-graders: 7%, 1^st^-graders: 37%), and 32% of all trials were removed (28% in 5^th^-graders, 29% in 3^rd^-graders, and 40% in 1^st^-graders). 2) in the remaining data, trials with incorrect and missed responses were discarded, as were trials with RTs below 200ms and above 5000ms, and above 2.5 standard deviations from individual participant’s mean RTs (per Gaspelin et al., 2015).

**Supplemental Results**

**Supplemental behavioural results**

Post-hoc t-tests on the main effect of Age revealed that 1^st^-graders were reliably slower than 3^rd^-graders (*t*_(33)_=3.8, *p* < 0.001), who were slower than 5^th^-graders (*t*_(32)_=4.9, *p* < 0.001).

**Supplemental N2pc results**

The 2 x 2 x 2 repeated-measures ANOVAs conducted on mean amplitudes from each age group’s electrode sites and time-windows revealed no significant main effects of Contralaterality, and therefore no N2pc, in any children (5^th^-graders (*F*_(1, 25)_*=*2.6, *p*=0.1), 3^rd^-graders (*F*_(1, 37)_*=*1.1, *p*=0.3), 1^st^-graders (*F*_(1, 27)_*=*0.2, *p*=0.7); Suppl. Fig. 1).

**Supplemental GFP results**

In 5^th^-graders, the main effect of Cue Colour, (*F*_(1, 25)_=0.8, *p*=0.4), and of Cue Modality (*F*_(1, 25)_*=*1.2, *p*=0.3), as well as the two-way interaction between these factors, (*F*_(1, 25)=_0.4, *p*=0.5), were not significant.

In 3^rd^-graders, the main effect of Cue Colour reached a nonsignificant trend (*F*_(1,37)_*=*3.07, *p*=0.09, η_p_²=0.08), while main effect of Cue Modality (*F*_(1, 37)_*=*1.5, *p*=0.2), and the two-way interaction between Cue Colour and Cue Modality were not significant (*F*_(1, 37)=_1.5, *p*=0.4).

Finally, in 1^st^-graders, the main effect of Cue Colour (*F*_(1, 27)=_0.3, *p*=0.6), and of Cue Modality (*F*_(1, 27)=_0.3, *p*=0.6), as well as the two-way interaction between these factors (*F*_(1, 27)=_0.08, *p*=0.7) were not significant.

**Supplemental ERP topography results**

In 5^th^-graders, the 3-way Map x Cue Colour x Cue Modality interaction, *F*_(8, 200)_*=*2.2, *p*=0.048, η_p_²=0.1, was first followed up as a function of Cue Colour. This revealed that target-colour cues, map durations between visual and audiovisual were comparable (all *p*’s > 0.1), while for nontarget-colour cues, the presence of Maps 52, 53, 54, and 59 differed. That is, Map53 was present longer for audiovisual (38ms) than visual cues (11ms), *t*_(25)_*=*4.2, *p=*0.005, and Map54 was present longer for audiovisual (30ms) than visual cues (11ms), *t*_(25)_*=*3.3, *p=*0.03. Meanwhile, Map52 was present shorter for audiovisual (4ms) than visual cues (23ms), *t*_(25)_*=*3.5, *p=*0.005, and Map59 was present shorter for audiovisual (8ms) than for visual cues (26ms), *t*_(25)_*=*3.1, *p=*0.04. Next, the interaction was followed up by Cue Modality. Here, for audiovisual cues, only Map54 was present longer for nontarget-colour cues (30ms) than for target-colour cues (6ms), *t*_(25)_*=*5.0, *p* < 0.001. For visual cues, Map51 was present longer for target-colour cues (24ms) than for nontarget-colour cues (3ms), *t*_(25)_*=*3.7, *p=*0.01, while Map59 was present shorter for target-colour cues (5ms) than for nontarget-colour cues (26ms), *t*_(25)_*=*3.1, *p=*0.007.

In 3^rd^-graders, the 3-way Map x Cue Colour x Cue Modality interaction, *F*_(3, 111.1)_*=*3.2, *p*=0.03, η_p_²=0.1, was first followed up as a function of Cue Colour. Here, for nontarget-colour cues, Map32 was present longer for audiovisual (56ms) than visual cues (25ms), *t*_(37)_=4.1, *p=*0.005, while Map31 and Map33 were both present shorter for audiovisual cues than visual cues (Map31: 27ms vs. 50ms, *t*_(37)_=3.1, *p=*0.004; Map33: 8ms vs. 21ms, *t*_(37)_=2.5, *p=*0.03). For target-colour cues, on the other hand, only Map34 was present longer for audiovisual (48ms) than for visual cues (18ms), *t*_(37)_*=*4, *p=*0.001. We next followed-up the interaction by Cue Modality. Here, for audiovisual cues, only Map32 was present shorter for target-colour cues (25ms) than nontarget-colour cues (56ms), *t*_(37)_=2.9, *p=*0.003. On the other hand, for visual cues, it was Map35 that was present longer for target-colour cues (30ms) than nontarget-colour cues (13ms), *t*_(37)_=3.2, *p=*0.005.

In 1^st^-graders, the 3-way Map x Cue Colour x Cue Modality interaction, *F*_(7, 189)_*=*2.2, *p*=0.04, η_p_²=0.1, was first followed up as a function of Cue Colour. For nontarget-colour cues, there were no significant differences between visual and audiovisual (all *p*’s > 0.1), but for target-colour cues, Map16 and Map18 were differentially present across visual and audiovisual V cues. Map16 was present longer for audiovisual (47ms) than visual cues (13ms), *t*_(27)_*=*2.8, *p=*0.02, while Map18 was present shorter for audiovisual (2ms) than visual cues (38ms), *t*_(27)_*=*3, *p=*0.007. Next we explored the interaction as a function of Cue Modality. For audiovisual cues, Map15 was present longer for target-colour cues (28ms) than nontarget-colour cues (4ms), *t*_(27)_=2.6, *p*=0.01. Meanwhile for visual cues, Map13 was present longer for target-colour cues (52ms) than nontarget-colour cues (6ms), *t*_(27)_=4, *p*=0.002, and Map18 was also present longer for target-colour cues (38ms) than nontarget-colour cues (12ms), *t*_(27)_=2.2, *p*=0.04. Conversely, Map14 was present shorter for target-colour cues (20ms) than for nontarget-colour cues (55ms) *t*_(27)_=2.9, *p*=0.05.

**Supplemental Discussion**

It is important to discuss in detail the fact that EN analyses - but not N2pc analyses - have both confirmed the evidence of visual attentional control found in behavior (two older groups) as well as revealed new findings that were not observed in behavior, such as evidence of visual control in the ERPs of 5-year-olds as well as of multisensory control in 7- and 9-year-olds. We will first discuss these in the context of the N2pc as a viable marker of attentional selection. Then, we will discuss the EN analyses in the context of their added benefits when studying brain mechanisms afforded by their multivariate nature.

**The viability of the N2pc as a marker of attentional selection**

The fact that the N2pc has not been observed in any of the developmental groups that we have tested deserves a careful discussion. We believe there are several likely factors that have contributed to the lack of N2pc’s observed in our study.

An intuitive explanation for the lack of recordable N2pcs in our study could be that the participating children did not pay attention as required by the task instructions. However, we know that this is not true, as all the groups, including the youngest group of 5-year-olds, performed way above the chance level (50%). Additionally, the two older groups showed behavioral evidence of at least visual control. Thus, poor attending is not a likely explanation for the current patterns of behavioral and brain results. Thus, the reasons for absence of the N2pc more likely lie in the specific brain response that it represents.

First, to date, the N2pc has been studied almost exclusively in the adults. In the rare exceptions where it was recorded in children, the youngest children in whom the N2pc was recorded were 10. However, we do not think that this means that young children cannot pay attention. We have already argued that children younger than 10 can pay attention selectively, even in the highly demanding attention task that we have used here. Thus, the N2pc is reliably observed from only a specific age.

Second, two studies so far reported N2pc in 10-year-olds (Couperus et al. 2015; Shimi et al. 2015). Even in those studies, where the N2pc was observed, it was recorded in conditions that we could be characterized as optimal for the recording of an attentional process. In the study of Couperus et al. (2015), the N2pc’s were recorded in response to targets in a circular search array; in the study of Shimi et al. (2015) – in response to targets that had to be remembered. In contrast, in our study, N2pc’s were recorded to distractors, rather than targets. This might have created conditions that were not optimal for observing N2pc, as the distractors were faint, equiluminant color changes. In line with this, the ERPs they elicited were small, compared to the ERPs to the targets. In conclusion, the N2pc requires optimal condition also in terms of physical and/or cognitive characteristics of the attended stimuli (e.g. they are target stimuli, for whom multiple, feature-, spatial, temporal and responses-related biases are supporting their processing).

Finally, and relatedly, in the two abovementioned studies, the children’s N2pc’s were slower to develop, slower to peak or transpired for longer, compared to the adults’ N2pc’s. This is a well-known characteristic of developmental counterparts of adult ERP components (at least in children older than infants). In this vein, the particular characteristics of our fast-paced task could have further added to the suboptimal conditions in which N2pc was to be observed. Namely, there was a window of app. 200-250ms between the onset of the distractor and the onset of the, larger, target-related ERPs. We have used that specific time-period in our previous studies, in adults, to prevent eye movements to the distractors, which would have eliminated any reliable ERP responses from being recorded. We have retained this exact time-window, knowing that children, especially the youngest ones, would be even more prone to moving their eyes. Thus, in conditions for the eliciting the N2pc, it is possible that one needs to create further accommodations in task and testing procedures, e.g., allowing sufficient time for the N2pc to develop fully in time in the course of the trial, which could compensate for the effects of the suboptimal conditions. Notably, these ‘optimal’ conditions would have a non-trivial effect on the behavior of and the cognitive processes elicited in the participants.

Thus, to summarize, the N2pc is not an automatic marker of attentional selection that is readily measurable whenever attention is studied with EEG/ERPs in a developmental context. Rather, it requires a wide range of conditions to be fulfilled in order to be readily observed, spanning participant characteristics (age), as well as physical and/or cognitive characteristics of the attended stimuli. In the case of such suboptimal conditions, accommodations, such as longer segments etc., may need to be necessary in order to record the N2pc. Overall, on the one hand, this suggests that there may exist a significant number of studies, in which such heightened conditions have not been fulfilled, and in which the N2pc has in fact not been recorded, where those results have not been published, thus leading to an erroneous view offered by the positive results that have been published, indicating that N2pc is an absolute brain response in any situations involving attentional selection of target objects from among nontarget objects (the so-called “file drawer problem”). On the other hand, these arguments make us even more assured as to our choice to use electrical neuroimaging to analyze ERP responses in our study. This is crucial as of central interest to us were the brain responses and mechanisms that supported attention (capture) to distractors that differed in their match with visual targets as well as in terms of the senses their engaged. Notably, we were already aware of the limitations of the N2pc when used as a marker of attentional selection (Matusz et al. 2019b; Turoman et al. 2021) when used in contexts outside of the traditional (arguably, the most optimal ones for eliciting the N2pc), visual paradigms. In contrast, electrical neuroimaging analyses have provided here the missing evidence for brain mechanisms supporting attentional capture (processes elicited between the cue and the onset of the target). Crucially, they did so while being robust against the conditions that were suboptimal for the recording of the N2pc: the age of the observer, the physical and cognitive characteristics of the stimuli to which the ERPs were recorded to, or the paradigm characteristics.

**The added benefits of electrical neuroimaging analyses when studying the brain function**

Our finding of task-set contingent attentional capture (our visual attentional control measure) in the behavior of two of the three child groups – 5^th^-graders and 3^rd^-graders – is valuable, as this cognitive process has until now been practically exclusively demonstrated in adults. The fact that evidence of modulations of ERPs by visual attentional control was found with EN analyses but not with N2pc provides evidence for heightened sensitivity of EN analyses over the N2pc analyses. By extension, the EN evidence of visual attentional control in the absence of behavioral evidence, in the 1^st^-graders, provides support for our EEG EN-derived results to be genuine. We take the systematicity of in comparing adults and children (reported in Turoman et al. 2021), and children across specific age groups a strong advantage of our approach when investigating cognitive and brain mechanisms governing developing visual and multisensory attentional control.

The presence of significant results obtained with EN analyses and absence of such with traditional N2pc analyses is well explained by the higher sensitivity of multivariate methods (EN analyses) as compared to univariate methods (canonical N2pc and ERP analyses). This sensitivity emerges from multivariate analyses capturing patterns in the data (such as topography x time patterns in the case of EEG) that the univariate analyses are not sensitive too. This sensitivity is well demonstrated in the scientific literature, including neuroscientific one, and is regarded as a fundamental mechanism by which the brain represents information (e.g., Haxby et al. 2001; Kriegeskorte et al. 2007; Habeck et al. 2008; Matusz et al. 2018).

This heightened sensitivity of multivariate methods compared to univariate methods is even more pertinent in the case of neurophysiological signals. Traditional ERP analyses involve analyzing data from just one electrode (or their small subset). This leads to the loss of signal, making univariate ERP analyses forcibly less sensitive. This approach also indicates a, more or less implicit, choice on the side of the experimenter that the brain activity measured at other sites on the scalp is irrelevant to the studied cognitive processes - in the case of the N2pc, attentional selection and its control. We have provided elsewhere an extensive discussion of the exact ways traditional ERP analyses are limited in conclusions they offer as to cognitive and brain mechanisms (Matusz et al. 2019b). Additionally, traditional ERP analyses are dependent on the choice of the reference. As such, the obtained results will be reproducible, across time or laboratories, only if the same reference is used, which makes them sensitive to the experimenters’ choices. Also, perhaps most importantly, those analyses rely on the fundamental assumption of fixed anatomy. That is, by recording the activity from always the same (one or several) electrode across all participants, it assumes that the underlying brain anatomy of all the participants’ brains is uniform, an assumption that is clearly difficult to defend.

Electrical neuroimaging analyses deal away with all those limitations. They are independent of the reference electrode, data-driven and consider data from the whole electrode montage. Crucially, as multivariate analyses, they tolerate well the variability in the underlying brain anatomy – exactly because they are sensitive patterns, here in spatiotemporal electrophysiological activity. If the organization of one particular set of anatomic generators in one participant is too different from the rest of the sample, a given map will simply not be fit to that person. However, the remaining brain activity, reliant on other brain networks, will be represented, by other maps. This all renders the spatiotemporal patterns in EEG by definition more reproducible than results of univariate EEG analyses.

Regarding the reproducibility of the specific topographic patterns produced by the EN analyses, the maps derived across similar paradigms have been extensively reproduced in the literature and match brain networks identified using hemodynamic methods. One example is template maps underlying EEG at rest, where a wide range of thought and brain activity can be expected. Across >10 studies from different research groups and using different rest protocols, a reliable number of 4-7 ‘resting state’ EEG maps have been demonstrated. Additionally, they have been reliably linked to specific resting-state brain networks identified using fMRI (reviewed in Michel & Koenig 2018).

**Supplemental References:**

Couperus, J. W., & Quirk, C. (2015). Visual search and the N2pc in children. *Attention, Perception, & Psychophysics, 77*, 768-776.

Eimer, M., Kiss, M., Press, C., & Sauter, D. (2009). The roles of feature-specific task set and bottom-up salience in attentional capture: an ERP study. *Journal of Experimental Psychology: Human Perception and Performance*, *35*(5), 1316.

Eimer, M., & Kiss, M. (2008). Involuntary attentional capture is determined by task set: Evidence from event-related brain potentials. *Journal of cognitive neuroscience*, *20*(8), 1423-1433.

Habeck, C., Foster, N. L., Perneczky, R., Kurz, A., Alexopoulos, P., Koeppe, R. A., ... & Stern, Y. (2008). Multivariate and univariate neuroimaging biomarkers of Alzheimer's disease. *Neuroimage*, 40, 1503-1515.

Haxby, J. V., Gobbini, M. I., Furey, M. L., Ishai, A., Schouten, J. L., & Pietrini, P. (2001). Distributed and overlapping representations of faces and objects in ventral temporal cortex. *Science, 293*, 2425-2430.

Kiss, M., Van Velzen, J., & Eimer, M. (2008). The N2pc component and its links to attention shifts and spatially selective visual processing. *Psychophysiology*, *45*(2), 240-249.

Luck, S. J., & Hillyard, S. A. (1994). Spatial filtering during visual search: evidence from human electrophysiology. *Journal of Experimental Psychology: Human Perception and Performance*, *20*(5), 1000.

Scerif, G., Kotsoni, E., & Casey, B. J. (2006). Functional neuroimaging of early cognitive development. *Handbook of Functional Neuroimaging of Cognition*, *351*.

Sun, M., Wang, E., Huang, J., Zhao, C., Guo, J., Li, D., ... & Song, Y. (2018). Attentional selection and suppression in children and adults. *Developmental science*, *21*(6), e12684.

**Supplementary Figure 1**

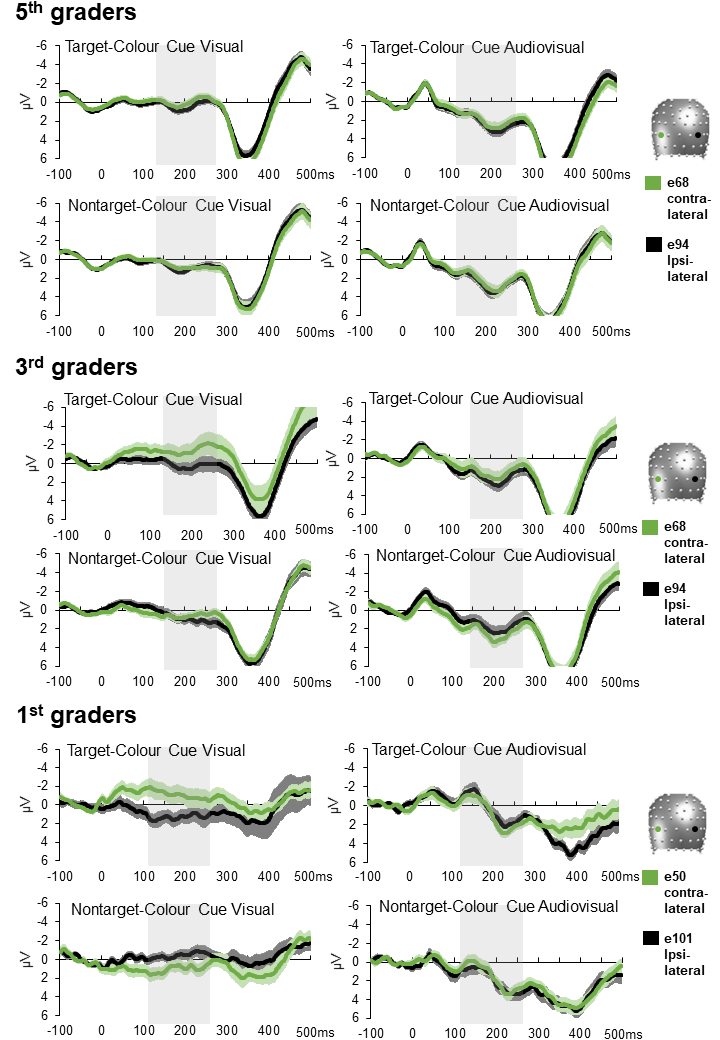

**Suppl. Fig. 1.** N2pc results showing mean amplitude values at contralateral and ipsilateral electrode sites, indicated in orange and black, per the legends on the figure. Each group’s own N2pc time-windows are highlighted in light grey. No groups show significant contra-ipsi differences, e.g. reliable N2pc’s.
